# DELE1 oligomerization promotes integrated stress response activation

**DOI:** 10.1101/2022.10.01.510468

**Authors:** Jie Yang, Kelsey R. Baron, Daniel E. Pride, Anette Schneemann, Xiaoyan Guo, Wenqian Chen, Albert S. Song, Giovanni Aviles, Martin Kampmann, R. Luke Wiseman, Gabriel C. Lander

## Abstract

Mitochondria are dynamic organelles that must continually adapt and respond to cellular stress. Recent studies demonstrated that mitochondrial stress can be relayed from mitochondria to the cytosol by the release of a C-terminal proteolytic fragment of DELE1 that binds to the eIF2α kinase HRI to initiate integrate stress response (ISR) signaling. Here, we report the cryo-electron microscopy structure of the active, C-terminal cleavage product of human DELE1 at *∼*3.8 A° resolution. Our structure reveals that DELE1 assembles into a high-order oligomer that is observed both *in vitro* and in mammalian cells. Structurally, the oligomer consists of eight DELE1 monomers that assemble with D4 symmetry via two sets of distinct hydrophobic inter-subunit interactions. We identified the key residues involved in DELE1 oligomerization, and confirmed their role in stabilizing the octamer *in vitro* and in cells using mutagenesis. Further, we show that assembly impaired DELE1 mutants are compromised in their ability to induce ISR activation in cell culture models. Together, our findings provide molecular insights into the activity of DELE1 and how it signals to promote ISR activity following mitochondrial insult.

## Introduction

Mitochondria are hubs of bioenergetic processes whose homeostasis must be tightly regulated for cellular function and longevity^1-4^. In response to physiological and pathological insults, mitochondria use molecular signals to initiate nuclear transcription of stress-response genes as a means of restoring cellular homeostasis. In mammals, mitochondrial stress is relayed to a central pathway known as the integrated stress response (ISR). Four kinases – HRI, PKR, PERK and GCN2 - serve as specific sensors for detecting distinct forms of cellular stress, and convergently phosphorylate a common substrate, eukaryotic initiation factor 2α (eIF2α)^5^. Phosphorylation of eIF2α leads to a global repression of protein production while increasing translation of specific proteins such as the transcription factor ATF4. The mitochondrially targeted protein DELE1 is a key signaling factor linking mitochondrial dysfunction to the ISR^6,7^. Mitochondrial stress activates the inner mitochondrial membrane protease OMA1, which promotes cleavage of DELE1, resulting in an accumulation of the DELE1 carboxy-terminal domain (DELE1^CTD^) in the cytosol, where it activates the HRI kinase. Although its critical role in relaying mitochondrial stress to activate the ISR is well-established^6-8^, we lack a molecular description of DELE1, which prevents us from understanding the mechanisms by which DELE1 mediates the activation and subsequent signaling of the ISR.

## Results

### DELE1^CTD^ cryo-EM structure reveals an oligomer

DELE1 is a 56 kiloDalton protein containing a mitochondrial targeting sequence and has been reported to be imported into mitochondria^6,7,9^. Although DELE1 has not been structurally characterized experimentally, the N-terminus is predicted by AlphaFold2 to be mostly unstructured, except for a hydrophobic α-helix. The C-terminus is predicted to consist of tetratricopeptide repeats (TPRs), comprising 11 α-helices (α1 – α11) (**Fig. 1a**). To gain a better understanding of how DELE1 functions in regulating ISR activity, we initially attempted to express and purify the full-length DELE1 and the active carboxy-terminal cleavage product of DELE (DELE1^CTD^) from either bacterial or mammalian expression systems, but both displayed low expression and poor solubility. However, expression and solubility of both constructs were improved by fusing a maltose-binding protein (MBP)^10^ to their N-terminus (**Fig. 1a**). Whereas the full-length MBP-DELE1 eluted from a size-exclusion chromatography (SEC) column in a peak volume of *∼* 15 ml, purified MBP-DELE1^CTD^ surprisingly eluted in a peak volume of *∼*11ml, corresponding to a much larger molecular weight, suggesting that this cleaved construct assembles into a higher-order oligomeric species (**Fig. 1b**).

**Figure 1.**
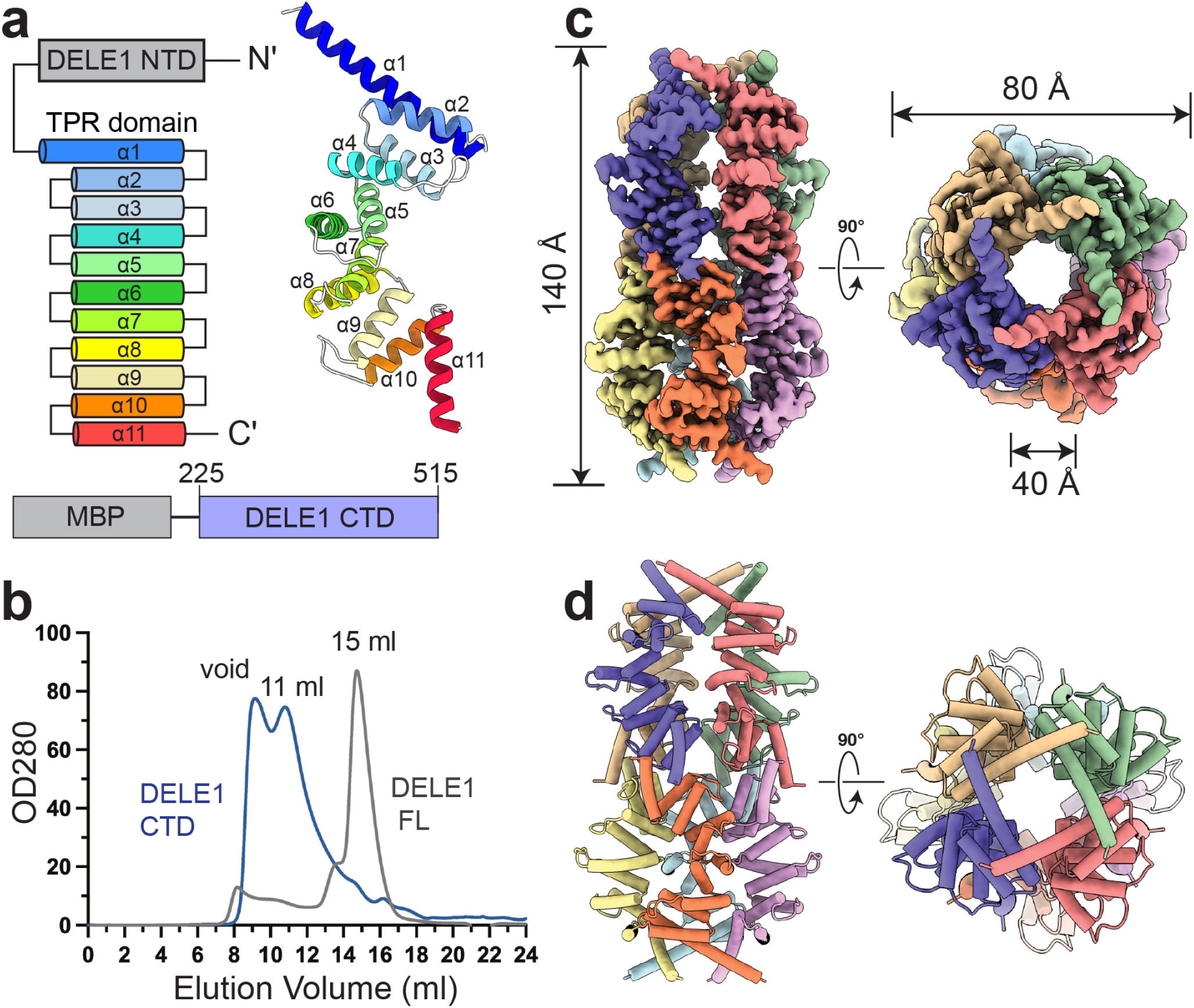
Cryo-EM structure of the oligomeric DELE1^CTD^. **(A)** Domain organization of the DELE1 protein with the TPR helices individually colored. To the right is a protomer from our cryo-EM structure, with the helices of the TPR domain colored as in the diagram. Below is a schematic of the MBP-tagged construct used for producing the DELE1 samples in this study. Residues 225-515 are designated as the DELE1 C-terminal domain (DELE1^CTD^). **(B)** Size-exclusion chromatograms of the purified recombinant MBP-tagged full-length DELE1 (gray), and DELE1 CTD (blue), which eluted at peak volumes of 15 and 11 ml, respectively. **(C)** Orthogonal views of the cryo-EM reconstruction of the D4-symmetric DELE1^CTD^ oligomer. Each subunit is denoted with a different color. **d**, The DELE1^CTD^ atomic model is shown using the same orientations and subunit coloring as in (C).

We next used single particle cryo-electron microscopy (cryo-EM) analysis to investigate the oligomeric structure of active DELE1^CTD^. Two-dimensional analysis of the cryo-EM data revealed that the MBP-DELE1^CTD^ assembles into an elongated, symmetric oligomer *∼*140 A° in length and *∼*80 A° in diameter (**Supplementary Fig. 1a, b**). Fuzzy densities observed at each end of the oligomer likely correspond to the flexible MBP solubilization tags, indicating that MBP-DELE1^CTD^ oligomerizes with N-termini at the periphery of the complex and C-termini directed toward the center (**Supplementary Fig. 1b**). Three-dimensional analyses confirmed that MBP-DELE1^CTD^ oligomerizes into a cylindrical D4-symmetric octamer that is mostly hollow and intrinsically flexible (**Fig. 1 c, d, Supplementary Fig. 1b**). Despite the flexibility of the oligomer, the reconstruction was resolved to a global resolution of*∼* 3.8 A°, which was sufficient to confidently build and refine an atomic model (**Supplementary Fig. 1b-e, Supplementary Table 1**). The MBP-DELE1^CTD^ octamer assembles as a dimer of two tetramers, where the N-termini from four monomers organize into a four-fold-symmetric crown-like assembly capping each end of the octamer. The TPRs from each monomer extend toward the center, giving the oligomer a spring-like appearance. Due to flexibility of the linker, the MBP moieties were not resolved in the reconstructed density, and thus do not impact the organization of the octamer.

### Two major interfaces mediate the DELE1 oligomer formation

The atomic model enabled us to closely examine the specific interactions involved in oligomerization of the DELE1^CTD^. Notably, there are only two sites of inter-protomer interactions involved in oligomerization of active DELE1^CTD^ - one at the N-terminal end (hereafter referred to as Interface I), and another near the C-terminus of the TPR (hereafter referred to as Interface II) (**Fig. 2a**). The tetrameric crown-like assembly at each end of the complex arises from interactions between the first helices (α1) of four DELE1^CTD^ protomers where F250 and L251 from one subunit are closely packed with L242 and L239 from the adjacent subunit (**Fig. 2b, d, Supplementary Fig. 2b**). These hydrophobic interactions, which have an estimated buried surface area of*∼* 518 A° ^2^ as calculated by the PISA server^11^ (**Supplementary Fig. 2a**), stabilize the four neighboring α1 helices into a weaved, square-like arrangement. We posit that the hydrophobic interactions within the α1 drives the oligomerization of DELE1 tetramers, and that assembly of the DELE1^CTD^ tetramer positions the C-terminal regions of the TPRs for octamer assembly. Our cryo-EM reconstruction shows that the C-terminal end of the TPRs engage in an interlocking network of interactions with a buried surface area of *∼*504 A° ^2^ (**Supplementary Fig. 2a**), which together stabilize the dimer of DELE1^CTD^ tetramers (**Fig. 2c**). The first of these interactions appears to be a strong hydrogen bond, based on the intensity of the cryo-EM density (**Supplementary Fig. 2d**), between R403 from the loop preceding α10 and S432 from α11 of the neighboring subunit (**Fig. 2e**). Next, residues L405 and L409 within α10 from adjacent subunits engage in symmetrically related hydrophobic interactions facing the lumen of the DELE1^CTD^ cylinder (**Fig. 2e, Supplementary Fig. 2e**). Lastly, the F431 residues from adjacent α11 subunits are positioned for a π-stacking interaction (**Fig. 2e, Supplementary Fig. 2c**). Together, these interactions form four interlocking “V-shaped” assemblies around the equator of the DELE1^CTD^ octamer.

**Figure 2.**
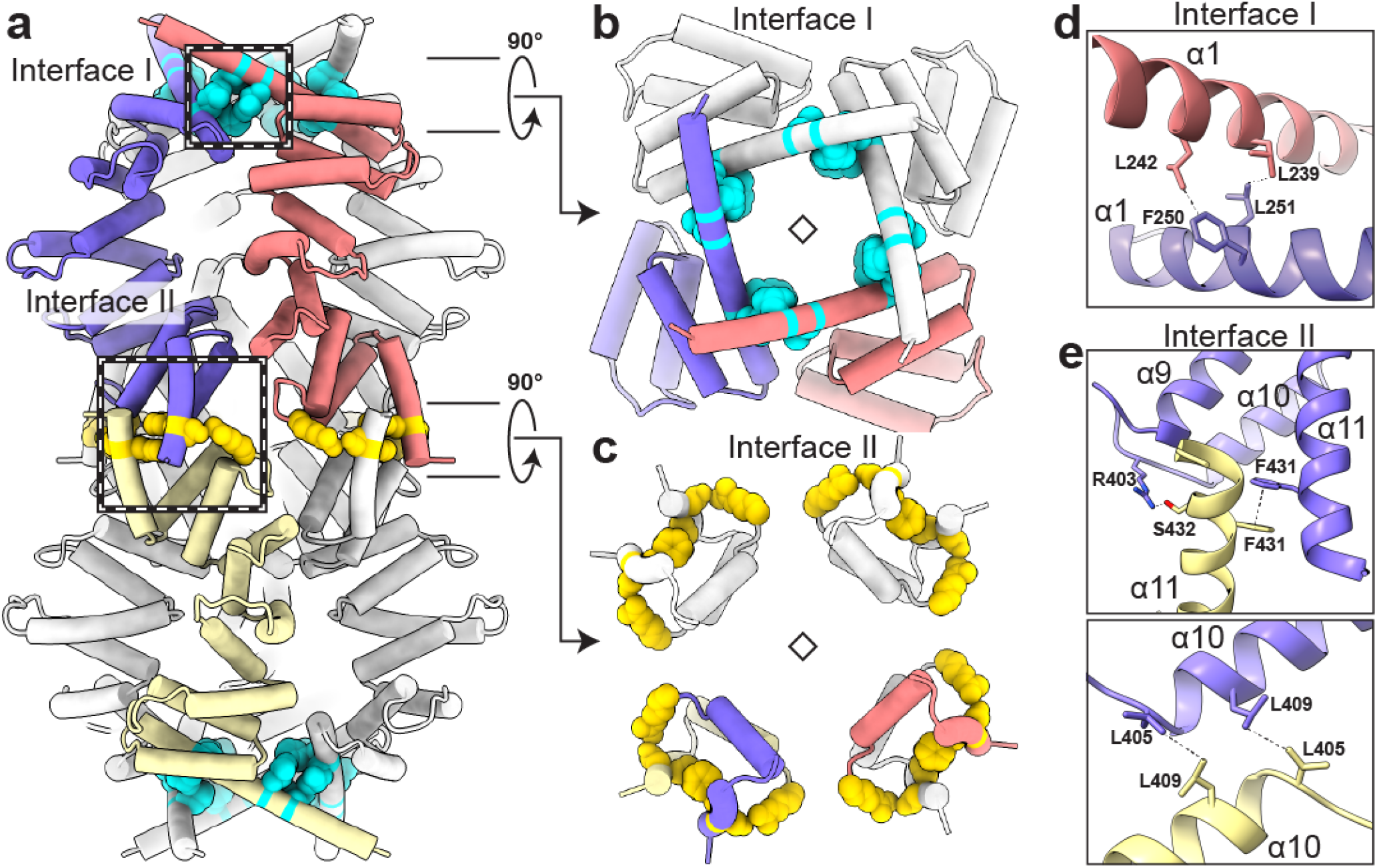
Two interfaces contribute to DELE1^CTD^ oligomerization. **(A)** A cartoon representation of the DELE1^CTD^ octamer, with three subunits highlighted to show the two distinct interaction interfaces, and residues involved in the interactions represented as spheres (cyan at Interface I, gold at Interface II). The purple subunit interacts with the red subunit at the N-terminal α1 crown, and interacts with yellow subunit at the C-terminus of the TPR. **(B)** An axial view of the DELE1^CTD^ crown, demonstrating how the inter-subunit interactions (cyan) link the four subunits together in a four-fold symmetric arrangement. **(C)** Axial view of a central cross-section of the DELE1^CTD^ octamer shows that the inter-subunit interactions (gold) establish dimeric assemblies. **(D)** A close-up view of the inter-protomer interactions at Interface I, demarcated by the upper dashed box in (A). Residues F250 and L251 in α1 from the purple-colored subunit interact with L239 and L242 in α1 of the neighboring red subunit. Hydrophobic interactions between F250/L2412 and L251/L239 are represented as dotted lines. **(E)** Top: close-up views of the subunit C-terminal TPR interface (Interface II), with interacting residues shown as sticks. R403 from the loop preceding α9 of the purple subunit hydrogen bonds with S432 in α11 of yellow subunit. Symmetrically related F431 residues from neighboring subunits are positioned for π-stacking interaction. Below: A close-up view of the inter-protomer interface visualized from the lumen, depicting the C2-symmetric interactions between L405 and L409 of the adjacent subunits.

### Structure-based mutations disrupt the DELE1 oligomer both *in vitro* and in cells

Having identified the specific residues involved in stabilization of the DELE1^CTD^ oligomer in our cryo-EM structure, we next sought to probe the biochemical and biophysical relevance of these interactions in DELE1 assembly using mutagenesis. We first aimed to disrupt the interactions at Interface II, which are involved in dimerization of DELE1^CTD^ tetramers according to our cryo-EM structure. This was achieved by introducing two mutations: R403A, which we anticipated would disrupt the strong inter-subunit hydrogen bonding we observed in the cryo-EM density, and F431S, which would abolish the observed inter-subunit π-stacking interactions. These mutations were cloned into the same MBP-DELE1^CTD^ fusion vector that was used for the wild type (WT) DELE1^CTD^. The SEC chromatogram of this double mutant confirmed our hypothesis that disrupting the interactions at Interface II would compromise octamer assembly, as we observed formation of an oligomeric species that was smaller than that observed for WT DELE1^CTD^ (**Fig. 3a)**. SDS-PAGE of the SEC peak fractions confirmed the presence of the R403A/F431S mutant protein **(Fig. 3h)**. We also used negative stain EM to confirm that the R403A/F431S double mutant prevents assembly of the octameric species, instead forming smaller oligomers that are consistent with DELE1^CTD^ tetramers (**Supplementary Fig. 3c, d**).

**Figure 3.**
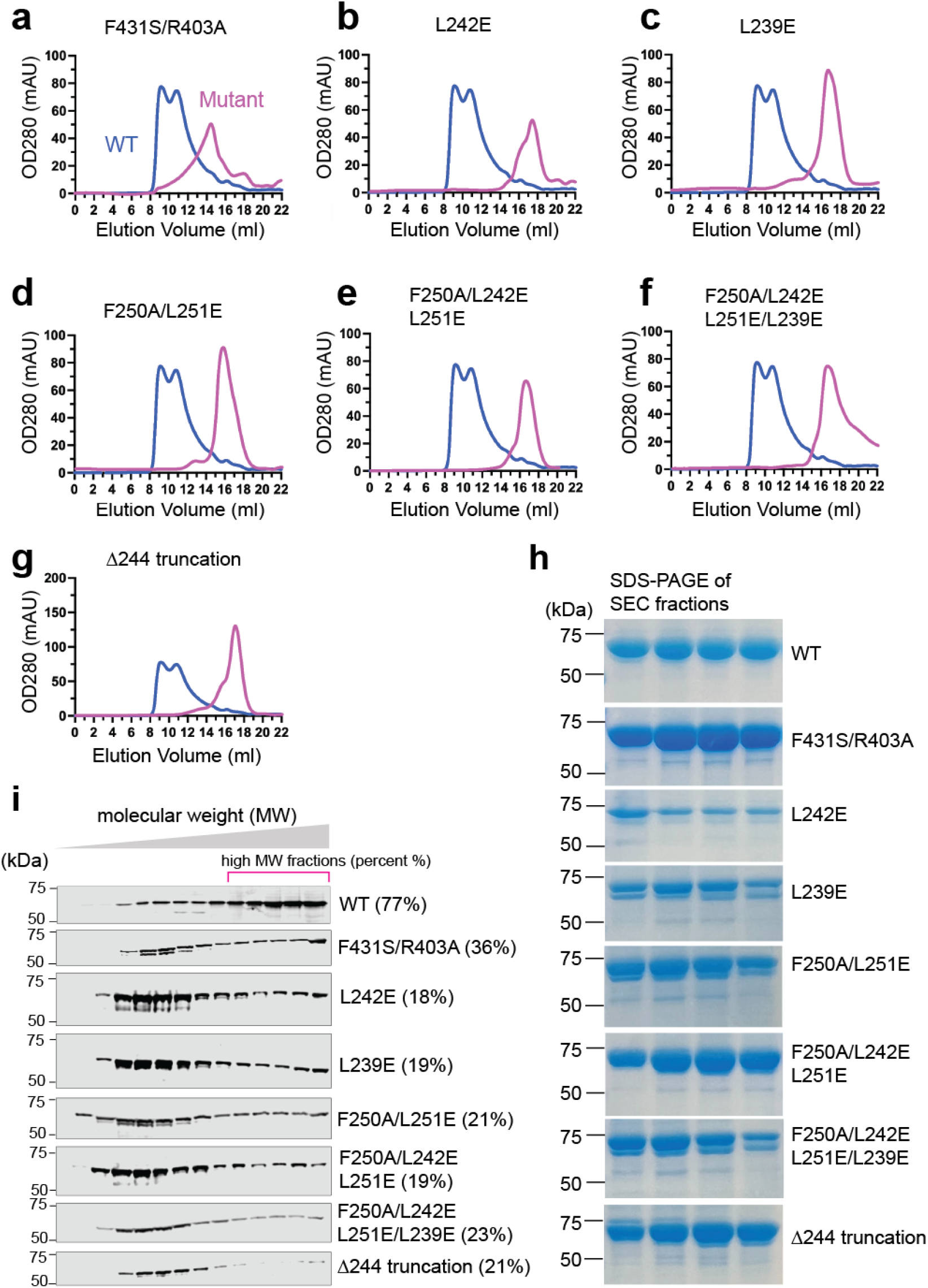
Structure-guided mutations disrupt DELE1^CTD^ oligomer formation. **(A)** Size-exclusion chromatograms of the WT DELE1^CTD^, and the Interface II mutation (labeled as R403A/F431S), which are colored blue, and magenta, respectively. **(B-G** Size-exclusion chromatograms of the WT DELE1^CTD^ overlaid with different Interface I mutations, including L242E (B), L239E (C), F250/L251E (D), L242E/F250A/L251E (E), L239E/L242E/F250A/L251E/ (F), and the 244 truncation (G). **(H)**, SDS-PAGE gels for the peak fractions of the WT and various DELE1^CTD^ mutants shown in (A-G). **(I)**, Immunoblot of WT and mutant DELE1^CTD^ proteins from sucrose gradient fractions. The 14 sucrose gradient fractions from light to heavy molecular weights (left to right) were blotted with an anti-GFP antibody to detect the DELE1^CTD^-GFP fusion proteins. The percent of the last five fractions were quantified and calculated against the total protein level in all fractions.

We next mutated the four residues within α1 that we identified as important for assembly of the tetrameric N-terminal crowns of the DELE1^CTD^ octamer. We also removed the entirety of α1 by truncating the N-terminus to V244 (Δ244). As shown in **Fig. 3b-g**, individual or combined mutations at the N-terminus, as well as truncation of α1, disrupted DELE1^CTD^ oligomerization, as demonstrated by SEC chromatograms of the mutant proteins corresponding to monomeric or dimeric species. The presence of mutants was validated by SDS-PAGE of the respective SEC peak fractions **(Fig. 3h)**. We further confirmed this abolishment of higher-order oligomerization by visualizing the mutant proteins with negative stain EM (**Supplementary Fig. 3e**). Together, these data support an assembly model wherein hydrophobic interactions at Interface I promote formation of the DELE1^CTD^ tetramers, and these tetramers dimerize via interactions at Interface II. Notably, none of these mutations affected the overall fold of the DELE1^CTD^ monomer, as predicted by AlphaFold2 (**Supplementary Fig. 3a, b**), indicating that while these mutations impact DELE1^CTD^ oligomerization, these disruptions are not likely due to the perturbed folding and stability of the protein.

Having demonstrated the oligomerization role of the indicated residues at Interfaces I and II *in vitro*, we next aimed to examine the impact of these mutations on DELE1^CTD^ oligomerization in mammalian cells. To accomplish this, we expressed WT DELE1^CTD^ and DELE1^CTD^ mutants in HEK293T cells and used sucrose gradient sedimentation to assess their oligomerization state in cell lysates. Given the lack of effective, commercially available antibodies to detect the DELE1 protein by immunoblotting^6^, we introduced a green fluorescent protein (GFP) tag at the C terminus of DELE1^CTD^ and used an anti-GFP antibody to detect the DELE-GFP fusion protein. Although the WT DELE1^CTD^ was detected across a wide range of the gradient fractions, the majority of the protein migrated to the high molecular weight fractions (**Fig. 3i**), confirming formation of DELE1^CTD^ oligomers in cells. The presence of smaller oligomeric species or monomers was expected, as we observed similar oligomeric heterogeneity for WT DELE1^CTD^ in our EM images (**Supplementary Fig. 1a**). While Interface II mutants (R403A/F431S) retained some oligomeric species (as predicted based on the tetramerization of this mutant *in vitro*; **Fig. 3a,i**), these mutants notably shifted the protein to lighter fractions of the gradient relative to the WT protein (**Fig. 3i**), consistent with our *in vitro* data (**Fig. 3a-3h, Supplementary Fig. 3c**,**d**). Taken together, these results indicate that the cleaved form of DELE1 assembles into oligomers, and that this oligomerization is mediated by localized interactions at two sites within the protein.

### Assembly-deficient DELE1^CTD^ mutants show impaired activation of the integrated stress response

Our data clearly show that DELE1^CTD^ has the capacity to form an oligomer, and we next sought to define the specific importance of DELE1 oligomerization in ISR activation. It was previously shown that overexpression of DELE1^CTD^-mClover induces HRI-dependent increases in the ISR-regulated protein ATF4^6^. Consistent with this, we observed increased ATF4 in HEK293T cells overexpressing DELE1^CTD^-mClover (**Fig. 4a, b**). Similarly, overexpression of DELE1^CTD^-mClover harboring R403A/F431S Interface II mutations, which abolished DELE1 octamer formation but maintained capacity to form tetramers (**Fig. 3a, i** and **Supplementary Fig. 3d**), led to an increase in ATF4 expression in HEK293T cells (red in **Fig. 4a, b**). However, overexpression of DELE1^CTD^ containing Interface I mutants, which completely disrupt DELE1 oligomerization *in vitro* and in cells (**Fig. 3b-g, i** and **Supplementary Fig. 3e**), did not lead to a notable increase in ATF4 protein levels (purple in **Fig. 4a, b**). Identical results were observed in HEK293T cells stably expressing an ATF4-FLuc ISR translational reporter (**Supplementary Fig. 4a**) transfected with DELE1^CTD^-GFP constructs containing Interface I mutations (**Fig. 4c**). The HRI activator BtdCPU^12^ was used as a control. Together, these results indicate that disruption of DELE1^CTD^ oligomerization prevents DELE1^CTD^-dependent ISR activation.

**Figure 4.**
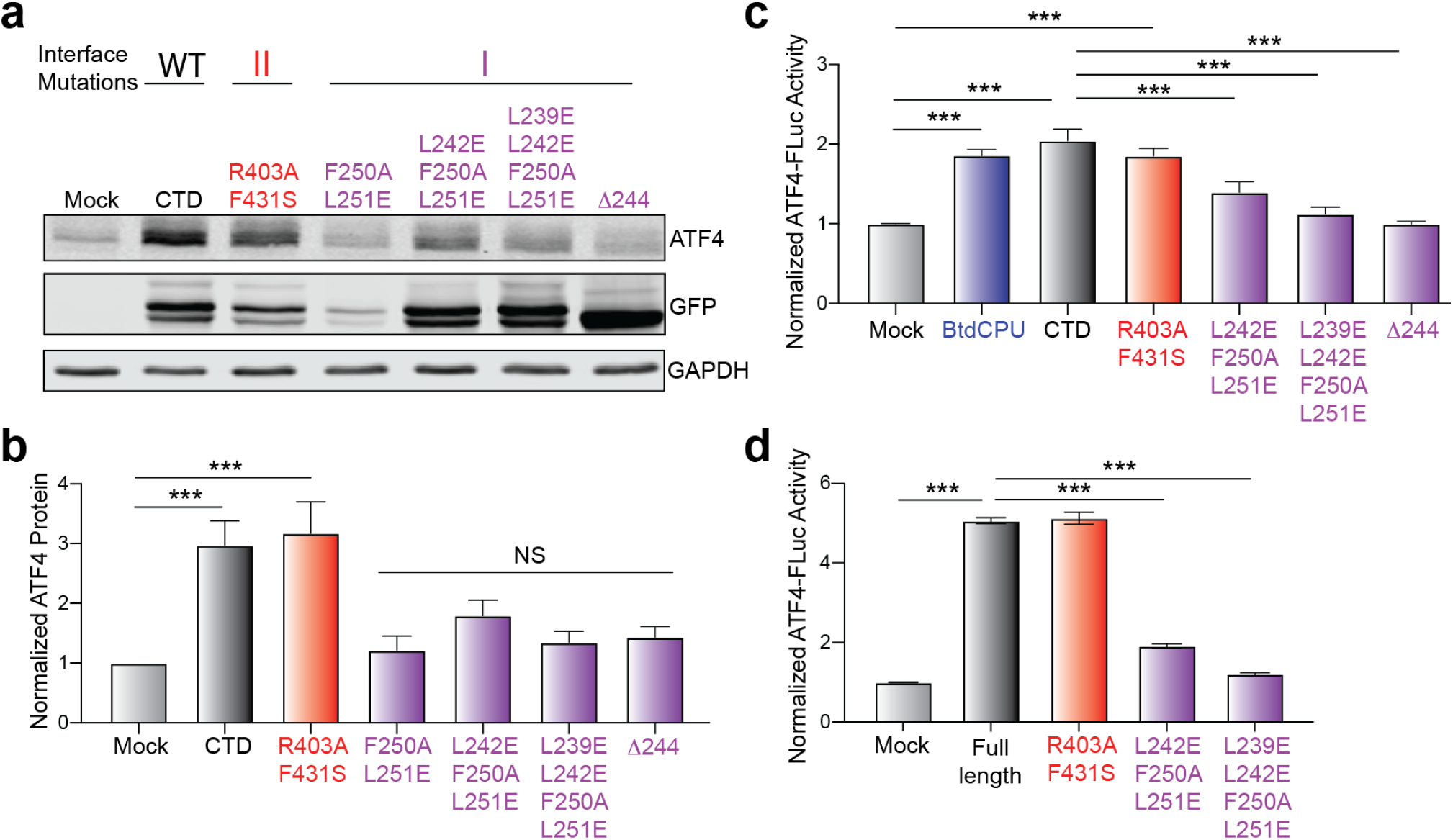
Disruption of DELE1 oligomerization abolishes ATF4 activation. All mutations are colored according to the DELE1 interface (purple for Interface I, red for Interface II). **(A)** An immunoblot of ATF4 levels at 24 hrs post-transfection with WT DELE^CTD^ (labeled as CTD), R403A/F431S mutant, F250A/L251E mutant, L242E/F250A/L251E, L239E/L242E/F250A/L251E, or the Δ244 truncation. **(B)** Quantification of the immunoblots in (A) (mean + s.d., n = 7 experimental repeats). **(C)** ATF4 activation measured by luciferase in cells, transfected with empty vector, treated with BtdCPU, or transfected with WT DELE^CTD^ (labeled as CTD), R403A/F431S mutant, F250A/L251E mutant, L242E/F250A/L251E, L239E/L242E/F250A/L251E, or the Δ244 truncation (mean + s.d., n = 6 experimental repeats). **(D)**The full-length WT DELE1, full-length R403A/F431S mutant, full-length L242E/F250A/L251E mutant, or full-length L239E/ L242E/F250A/L251E/ mutant were expressed in the DELE1 KD HEK293T cells. The ATF4 reporter level were measured 16 hrs using flow cytometry.

Previous results showed that DELE1^CTD^ binds to HRI to induce its activation^6,7^. Thus, we next sought to define whether HRI binding is dependent on DELE1^CTD^ oligomerization. We co-overexpressed FLAG-HRI and DELE1^CTD^-GFP mutants in HEK293T cells and monitored the recovery of DELE1^CTD^ in FLAG-HRI immunopurifications (IPs). Surprisingly, all DELE1^CTD^ mutants efficiently co-immunopurified with FLAG-HRI (**Supplementary Fig. 4b**), indicating that DELE1 oligomerization is not required for interaction with HRI.

Apart from DELE1^CTD^, recent results showed that full-length DELE1 can similarly accumulate in the cytosol and activate HRI-dependent ISR signaling in response to specific types of stress^9,13^. This observation suggests that full-length DELE1 should similarly assemble into oligomers to promote ISR activation. Consistent with this, we found that both full-length and C-terminally truncated DELE1-mClover migrated as oligomers in sucrose gradients of lysates prepared from HEK293T cells transiently transfected with full-length DELE1 (**Supplementary Fig. 4c**), indicating that full-length, mature DELE1 can integrate into oligomers. Further, we found that overexpression of full-length DELE1 robustly activated the ATF4-FLuc ISR reporter in HEK293T cells (**Fig. 4d**). Similar activity was observed for full-length DELE1 harboring the Interface II mutations R403A and F431S. However, Interface I mutants prevented ATF4-FLuc activation afforded by DELE1 overexpression. Collectively, these results indicate that both full-length DELE1 and DELE1^CTD^ can integrate into oligomers to initiate ISR signaling, and their ability to induce ISR is compromised by oligomerization-deficient mutants.

## Discussion

DELE1 was previously identified to function as a messenger protein relaying mitochondrial stress to the ISR, enabling a robust and precise cellular response to various mitochondrial perturbations. Our studies suggest that cytosolic DELE1 assembles as an oligomer that is required to initiate downstream ISR signaling. Given the bioenergetic demands of mitochondria and their dynamic relationship to cellular function, DELE1-depenent HRI activation must be tightly controlled to prevent activation under resting conditions, but also allow rapid triggering to counter insults associated with physiopathological stresses^2,14-17^. Our findings reveal a molecular mechanism that enables the precise regulation of the DELE1-mediated stress signaling under diverse cellular conditions, wherein DELE1 oligomerization serves as a molecular signal to activate the ISR.

The importance of DELE1 oligomerization suggests new potential mechanisms to distinctly regulate ISR signaling through this pathway in response to diverse types of pathologic insults. Post-translational DELE1 modification or altered interaction with cytosolic factors that influence DELE1 oligomer stability could directly promote or suppress ISR activity. Further, while full-length DELE1 and C-terminal cleaved DELE1^CTD^ both incorporate into oligomers, differences in the stability or regulation of these distinct oligomer populations could enable adjustments in the extents and durations of DELE1-mediated ISR signaling in response to the mitochondrial insults. Consistent with this, we show that fulllength DELE1 does not readily assemble into oligomers *in vitro* (**Fig. 1b**), suggesting that this protein may less effectively oligomerize in response to mitochondrial stress. While the potential to regulate DELE1-dependent ISR signaling through post-translational regulation of DELE1 oligomerization remains to be further explored, our findings suggest that modulation of ISR signaling *via* DELE1 oligomerization enables tuning of the DELE1-mediated response to varying levels and types of cellular stress.

Prior studies showed that cleaved DELE1 directly binds HRI for activation, but the molecular mechanism of DELE1-induced HRI activation is unknown. Here we show that WT and oligomerization-deficient DELE1 mutants all have the capacity to bind HRI, but oligomerization-deficient DELE1 mutants abrogate ISR activation. This suggests that DELE1 functions as a template for HRI dimerization and subsequent activation. Other ISR-associated kinases such as PKR undergo ligand-induced dimerization for trans-autophosphorylation and enzymatic activation^18,19^. It is plausible that HRI may use a similar activation mechanism, whereby DELE1^CTD^ oligomerization produces a signaling platform that scaffolds HRI for activation. Further investigation into the details of the DELE1-HRI interaction and how these interactions mediate HRI kinase activation will establish a more complete mechanistic understandings of molecular signaling linking mitochondrial stress to the ISR.

Rapid and efficient transduction of mitochondrial stress to the ISR is an essential component of cellular homeostasis maintenance, and dysregulation of this pathway has been observed in a wide range of human diseases^14-16,20^. It has proven particularly challenging to target mitochondrial function in the context of pathological conditions due to the double-membrane barrier of the organelle and its dynamic morphology. DELE1 functions as a bridging effector of mitochondrial stress with access to the cytosol, positioning it as an attractive target for therapeutic purposes. Our description of the DELE1^CTD^ oligomer and its relevance to the ISR pathway provides a potentially serendipitous and novel avenue to specifically target DELE1 activity and subsequently tune adaptive ISR signaling for therapeutic intervention to counter human mitochondrial pathologies without impacting cellular capacity of responding to other stresses such as ER stress or viral infections.

## Materials and Methods

### Reagents, plasmids and cell lines

BtdCPU compounds were purchased from Fisher Scientific. The DELE1 cDNA was from GenScript. The anti-eGFP monoclonal antibody (F56-6A1.2.3) was purchased from Abcam. The anti-ATF4 monoclonal antibody was purchased from Cell Signaling (D4B8). The anti-GAPDH monoclonal antibody (sc-47724) was from Santa Cruz Biotechnology. The secondary antibodies against mouse and rabbit IgG were purchased from LI-COR Biosciences and all antibodies were diluted according to the manufacturer’s instructions. Gibson assembly method (NEB E2611) was used for all cloning experiments. The Q5 site-directed mutagenesis kit from New England Biolabs was used to generate the DELE1 mutants. The DELE1 full length, CTD and mutants for recombinant protein expressions were cloned into a modified pET22b bacterial expression vector, which was shared by Dr. Tsan Sam Xiao (Case Western Reserve University, Cleveland). The vector contains a non-cleavage maltose binding protein (MBP) tag at the N terminus and 6x-His tag at the C-terminus. For mammalian expression, DELE1 constructs were cloned into a modified pcDNA3.1 vector which contains a GFP tag at the C terminus. A six-residue Gly-Ser linker was engineered between DELE1 and the GFP tag. The ATF4-luciferase translational reporter cell line (ATF4-FLuc) was generated through the adaption of the previously published ATF4-mAPPLE reporter cell line6. Briefly, we replaced the mApple with firefly luciferase and stably transfected this reporter into HEK293T cells. Next, we selected a clonal cell line that showed robust activation of ATF4-Fluc upon treatment with various ISR activators, including oliomycin A (OA) and thapsigargin (Tg), demonstrating that the reporter cells can accurately report on ISR activation induced by multiple stressors.

### Protein expression and purification

The DELE1 full length, CTD, and mutants were all expressed and purified using a similar methodology. E. coli strain BL21 (DE3) Codon Plus RIPL (Agilent Technologies) was used for recombinant DELE1 protein expression. The transformed BL21 RIPL cells were grown at 37◦C until the OD reached ∼0.8 and protein expression was induced with 1 mM IPTG at 16°C overnight. Cells were collected by centrifuging at 5,000 x g for 30 minutes at 4 ◦C. The cell pellets of the DELE1 CTD and mutants were resuspended and lysed using sonication in a Tris lysis buffer (pH 8.0) containing and lysed using sonication. The lysate was cleared by ultracentrifuging at 30,000 x g for 30 minutes at 4 ◦C, followed by Ni-NTA affinity binding. The column was washed with lysis buffer supplemented with 25mM imidazole (10 column volumes) and 50mM imidazole (5 column volumes), respectively. Right after washing, the protein was eluted with elution buffer containing lysis buffer supplemented with 300mM imidazole. The full-length DELE1 was resuspended and sonicated in the same lysis buffer, and then solubilized and extracted from membranes by adding 1% DDM detergent while stirring at4 ^*◦*^C for 1 hr. The lysate containing solubilized DELE1 was clarified by ultracentrifuge at 45,000 x g for 1 hr at 4 ^*◦*^C, followed by Ni-NTA affinity binding. The column was washed with lysis buffer supplemented with 0.1% DDM and 25mM imidazole (10 column volumes), and eluted with lysis buffer supplemented with 0.1% DDM and 300mM imidazole. All proteins were further purified through SEC using a Superose 6 increase column. Peak fractions containing the target proteins were pooled, concentrated, aliquoted, and flash-frozen in liquid nitrogen for storage at −80 ^*◦*^C.

### Sample preparation for electron microscopy

For negative stain electron microscopy, 4 µl of DELE1 samples at 0.1 mg/ml were applied onto carbon coated 400 A° mesh Cu-Rh maxtaform grids (Electron Microscopy Sciences) which were plasma cleaned for 15 s using Pelco glow discharge cleaning system (Ted Pella, Inc.) with atmospheric gases at 15 mA. The samples were stained using 2% (w/v) uranyl formate solution, then blotted with Whatman #1 filter paper and dried. For cryo-EM, 4 µl of DELE WT CTD sample at 3 mg/ml was applied onto 300 mesh R1.2/1.3 UltraAuFoil holey gold grids (Quantifoil) that were plasma cleaned for 15 s using Pelco glow discharge cleaning system (Ted Pella, Inc.) with atmospheric gases at 15 mA. The grids were immediately vitrified by plunge-freezing using a Thermo Fisher Vitrobot Mark IV at 4 ^*◦*^C, 5 s, 100% humidity, and a blot force of 0 using Whatman #1 filter paper.

### Cryo-EM data collection and processing

Single particle cryo-EM data were collected on a Thermo-Fisher Talos Arctica transmission electron microscope operating at 200 keV using parallel illumination conditions^21,22^. Micrographs were acquired using a Gatan K2 Summit direct electron detector with a total electron exposure of 50 e^-^/A° ^2^ as a 49-frame dose-fractionated movie during a 9800-ms exposure time. The Leginon data collection software^23^ was used to collect a total of 13,164 micrographs during multiple sessions at 36,000 X magnification (1.15 A° /pixel) with a nominal defocus ranging from −0.7 µm to −1.2 µm. Stage movement was used to target the center of either four or sixteen 1.2 µm holes for focusing, and image shift was used to acquire high magnification images.

The Appion image processing wrapper^24^ was used to run MotionCor2 for real-time micrograph frame alignment and dose-weighting during data collection^25^. All subsequent image processing was performed in cryoSPARC^26^. The contrast transfer function (CTF) of each image was estimated using Patch CTF estimation (multi) using default parameters. Blob picking on a small portion of the CTF estimated images (*∼*200 images) using a particle diameter of 150 A° to 250 A° was used for initial particle picking. The picked particle coordinates were extracted at 1.15 A° /pixel from the motion-corrected and dose-weighted micrographs using a box size of 256 pixels, followed by 2D classification into 50 classes using default parameters. Representative views of 2D classes from the initial processing were used as templates for template-based particle picking in cryoSPARC, which resulted in *∼*1.06 million particle selections. After inspecting the particle picks, 660,425 picked particle coordinates with NCC score above 0.180, and local power between 85 and 166 were extracted at 1.15 A° /pixel from the motion-corrected and dose-weighted micrographs using a box size of 256 pixels. The extracted particles were two-dimensionally classified into 100 classes using default cryoSPARC parameters and the 92,455 particles contributing to the 2D classes displaying high-resolution secondary structural features were selected for further processing. A reference-free 3D ab-initio model was generated using default parameters and used for non-uniform refinement with C1 symmetry. The resulting reconstruction, which was resolved to 5.4 resolution according to the Fourier Shell Correlation (FSC) at a cutoff of 0.143, exhibited secondary structural features that were consistent with a D4-symmetry. Another round of non-uniform refinement was performed using the same Ab-initio but with D4 symmetry imposed. Heterogeneous refinement with D4 symmetry was then used to classify the particles into three classes, and the 67,067 particles contributing to the two classes containing higher resolution features were combined and then subjected to a final round of non-uniform refinement with D4 symmetry. This final refinement produced the final reconstruction at a reported global resolution of 3.8 A° at an FSC cutoff of 0.143. CTF refinement in cryoSPARC did not improve the resolution of the reconstruction. Directional FSCs were estimated by the 3DFSC server^27^ (**Supplementary Fig. 1**).

### Atomic model building and refinement

The AlphaFold2 model of DELE1 (**Supplementary Fig. 3a**) was initially docked into the reconstructed density map using UCSF Chimera. Real-space structural refinement and manual model building was performed with Phenix^28^ and Coot^29^, respectively. Phenix real-space refinement included global minimization, rigid body, local grid search, atomic displacement parameters, and morphing for the first cycle. It was run for 100 iterations, five macro cycles, with a target bonds RMSD of 0.01 and a target angles RMSD of 1.0. The refinement settings also included secondary structure and Ramachandran restraints. Figures for publication were prepared using UCSF Chimera^28^ and UCSF ChimeraX^30,31^. Data collection, refinement, and validation statistics are summarized in **Supplementary Table 1**.

### Sucrose gradient sedimentation

HEK293T cells at approximately 50% confluency on a 10 cm dish were transfected with 5 µg of DELE1 plasmid using calcium phosphate method, and the proteins were expressed for 24 hrs post-transfection. The cells were pelleted at 600 x g for 5 minutes at 4 ^*◦*^C and washed twice with 10 ml of PBS buffer. The final cell pellet was resuspended in 250 µl of lysis buffer (10 mM Tris/HCl pH 7.5, 150 mM NaCl, 0.5 mM EDTA, 0.5% (v/v) NP-40, 5% (v/v) glycerol, Roche complete protease inhibitors) and incubated on ice for 25 minutes. The lysate was then clarified by centrifugation at 21,000x g for 10 minutes at 4 ^*◦*^C and added on top of a 4 ml continuous 10-30% (wt/wt) sucrose gradient in TBS (25mM Tris-HCl/150mM NaCl, pH7.2). This was followed by centrifugation in a SW60Ti rotor at 33,000 rpm (146,334x g) for 18 hrs at 4 ^*◦*^C. The gradient was manually fractionated from the top into 0.2 ml fractions. The presence of DELE1 proteins in sucrose fractions was examined by immunoblotting using an anti-GFP antibody.

### Luciferase assay

ATF4-Fluc cells were transfected with 5 µg of each DNA using calcium phosphate. The media was replaced with fresh media the following morning after transfection. 16 hrs posttransfection, cells were seeded at 15,000 cells per well into both white and clear 96-well plates for luciferase and viability assays. 40 hrs post-transfection, mock transfected cells were treated with DMSO or 10 µM BtdCPU for eight hrs. Following drug treatment, Bright-Glo reagent was added to each well at a 1:1 ratio with respect to media volume. Cells were lysed for 10 mins at room temperate and luminescence signal was read on a Tecan machine with an integration time of 1000 ms.

### DELE1 rescue experiments in KD cells

Different DELE1 variants fused with GFP were transfected using Mirus TransIT ® -LT1 Transfection Reagent (MIR2304) into ATF4 reporter cells with about 50% of cell population having DELE1 KD. 24 hrs post transfection, cells were seeded in 96 well plate at about 25% confluency. The following day, oligomycin at a final concentration of 1.25 ng/mL was added for 16 hrs. After treatment, the cells were sorted into four different populations including DELE KD(+) GFP(+), DELE KD(+) GFP(-), DELE1 KD(-) GFP(+), DELE1 KD(-) GFP(-). ATF4 reporter level for each population of cells was measured using flow cytometry.

### Co-immunoprecipitation

Co-IP assay was performed to analyze the interaction between HRI and DELE1. HEK 293T cells were transiently transfected with 5 µg of FLAG tagged full-length HRI construct and 5 µg of GFP tagged DELE1 constructs (DELE1^CTD^ or mutants) using Lipofectamine 3000 transfection reagent (L3000008, ThermoFisher Scientific). GFP empty vector was used as a negative control. Cells were collected 24 hrs after transfection and lysed using a lysis buffer (25mM Tris/Cl pH 7.5; 150mM NaCl; 0.5M EDTA; 0.5% NP-40, 2.5% Glycerol, Roche complete protease inhibitors). The lysates were then incubated with 15 µl anti-FLAG M2 magnetic beads (M8823, Sigma-Aldrich) for three hours at 4 ^*◦*^C. Samples were washed three times with wash buffer (25 mM Tris-HCl, 150 mM NaCl, 0.1% NP-40). Proteins captured on the magnetic beads were eluted with 2X SDS loading dye for 5 min and analyzed by Western Blotting using rabbit anti-FLAG (14793S; Cell Signaling Technology) and anti-GFP (A-6455, ThermoFisher Scientific) to detect HRI-FLAG and DELE1-GFP, respectively.

## Data availability

The cryo-EM reconstruction and associated atomic model of DELE1^CTD^ have been deposited to the Electron Microscopy Data Bank (EMDB) and Protein Data Bank (PDB) under accession numbers EMDB: EMD-27269 and PDB: 8D9X, respectively. All data needed to evaluate the conclusions in the paper are present in the paper and the supporting information.

## Acknowledgements

We thank Bill Anderson at the Scripps Research Electron Microscopy Facility for microscopy support. We thank Charles Bowman and Jean-Christophe Ducom at Scripps Research High Performance Computing core for computational support. We thank Dr. Tsan Sam Xiao at Case Western Reserve University for kindly sharing the MBP-tagged vectors. We thank Dr. Benjamin Basanta for his valuable comments on designing DELE1 mutations. This work is supported by the National Institutes of Health (NIH) (NS095892 to R.L.W. and G.C.L, and NS125674 to R.L.W.). A.S.S. is supported by the NIH F31AG071162 and the Olson-King Endowed Skaggs Fellowship from Scripps Research. Computational analyses of EM data were performed using shared instrumentation funded by NIH S10OD021634 to G.C.L.

## Author contributions

J.Y. and G.C.L. conceptualized and led the project. J.Y. created all the constructs, purified the proteins, performed all cryo-EM structure determination, model building and refinement, and mechanistic interpretation. K.B. and R.L.W. designed and performed the ATF4 immunoblotting and ATF4 luciferase assay. D.P. assisted in purifying the DELE1 mutant recombinant proteins. A.S. performed the sucrose gradient sedimentation assay. X.G. performed the DELE1 rescue experiment. J.Y. and W.C. performed the Co-IP assay. A.S.S. assisted in the western blot. X.G., G.A. and M.K. generated the ATF4 Fluc reporter cell line. X.G. performed the fulllength DELE1 rescue assay. J.Y. wrote the original draft of the manuscript. G.C.L. and R.L.W. reviewed and edited the manuscript. All authors commented on the manuscript.

## Supplementary Materials

**Supplementary Figure 1.**
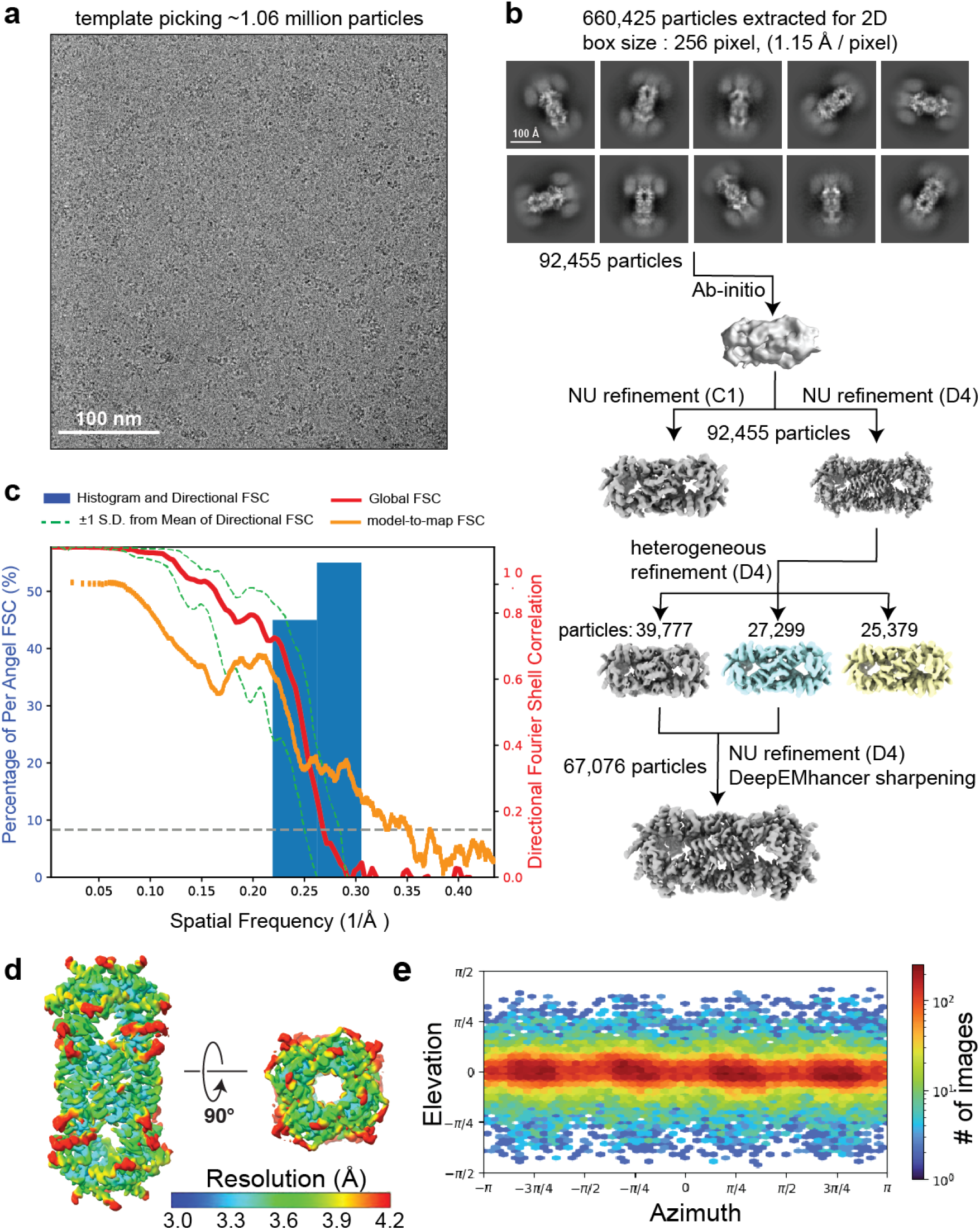
Cryo-EM structure determination of DELE1. **(A)** Representative micrograph of cryo-EM data collection. The high background on the micrograph appears to be the monomeric or small oligomeric species of DELE1^CTD^, possibly caused by denaturing due to interaction with the air-water interface during cryo-freezing^32,33^, although there may be oligomeric heterogeneity within the sample prior to freezing. **(B)** Cryo-EM data processing workflow using cryoSPARC software. The final 3D reconstruction map was used for model building and refinement. **(C)** 3D Fourier Shell Correlation (3DFSC)^27^ of the final reconstruction reporting a global resolution at *∼*3.8 A° and high level of resolution isotropy, with the FSC at 0.143 denoted. The model-to-map FSC plot from Phenix real space refinement is overlaid. **(D)** Final reconstruction filtered and colored by local resolution, as estimated in cryoSPARC. **(E)** Euler angle distribution plot of the particles used in the final 3D reconstruction showing complete tomographic coverage of projections.

**Supplementary Figure 2.**
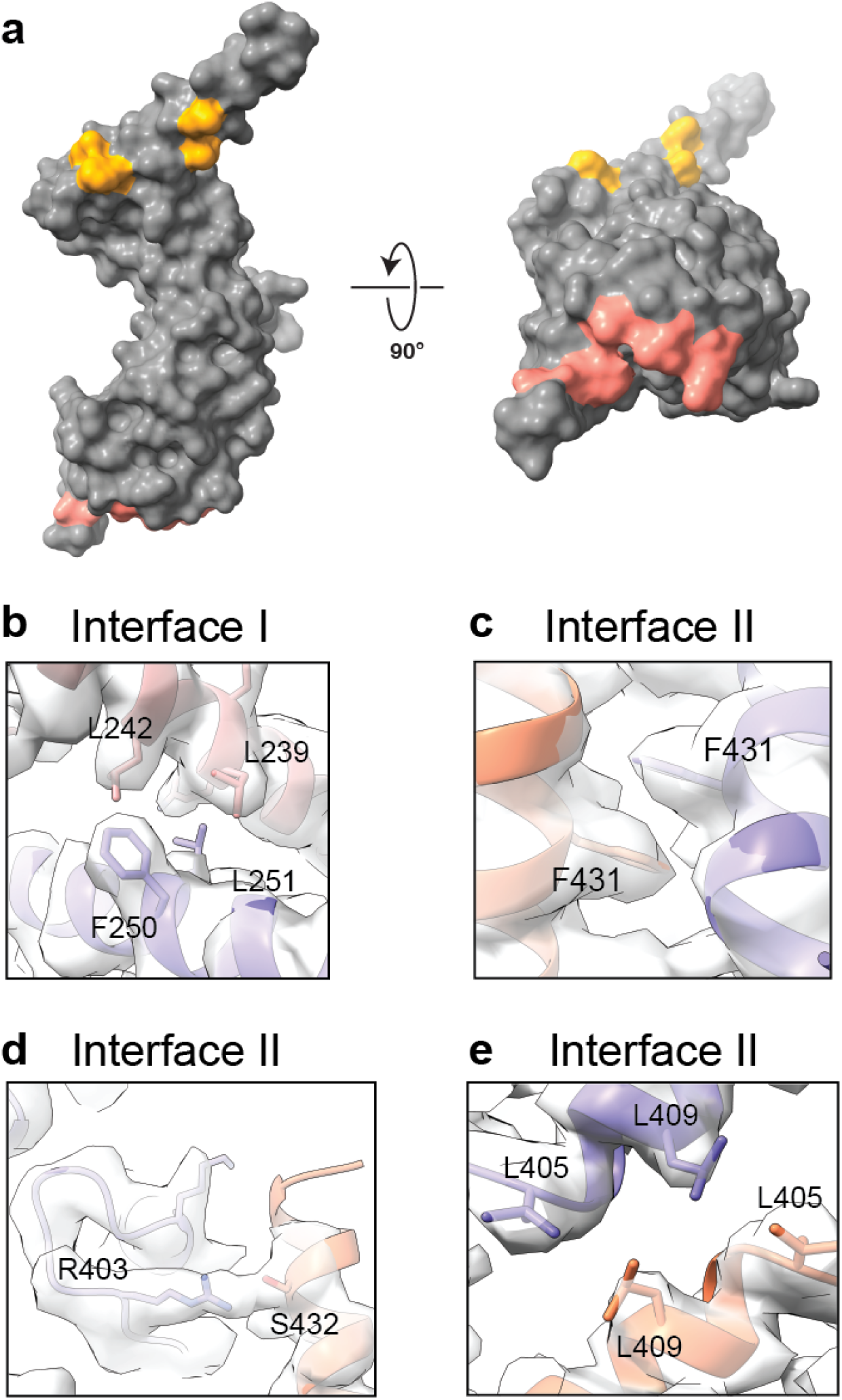
Inter-subunit interactions among the DELE1 oligomer. **(A)** A surface representation of the DELE1^CTD^ monomer is shown in two orthogonal views, colored gray. The buried surface areas associated with oligomerization Interfaces I and II are colored orange and salmon, respectively. **(B-E)** Detailed views of the Interface I and II interactions demonstrating the quality of the cryo-EM density, which is shown as transparent surface. The interacting residues are shown as sticks, and the coloring is consistent with that of **Fig. 2**.

**Supplementary Figure 3.**
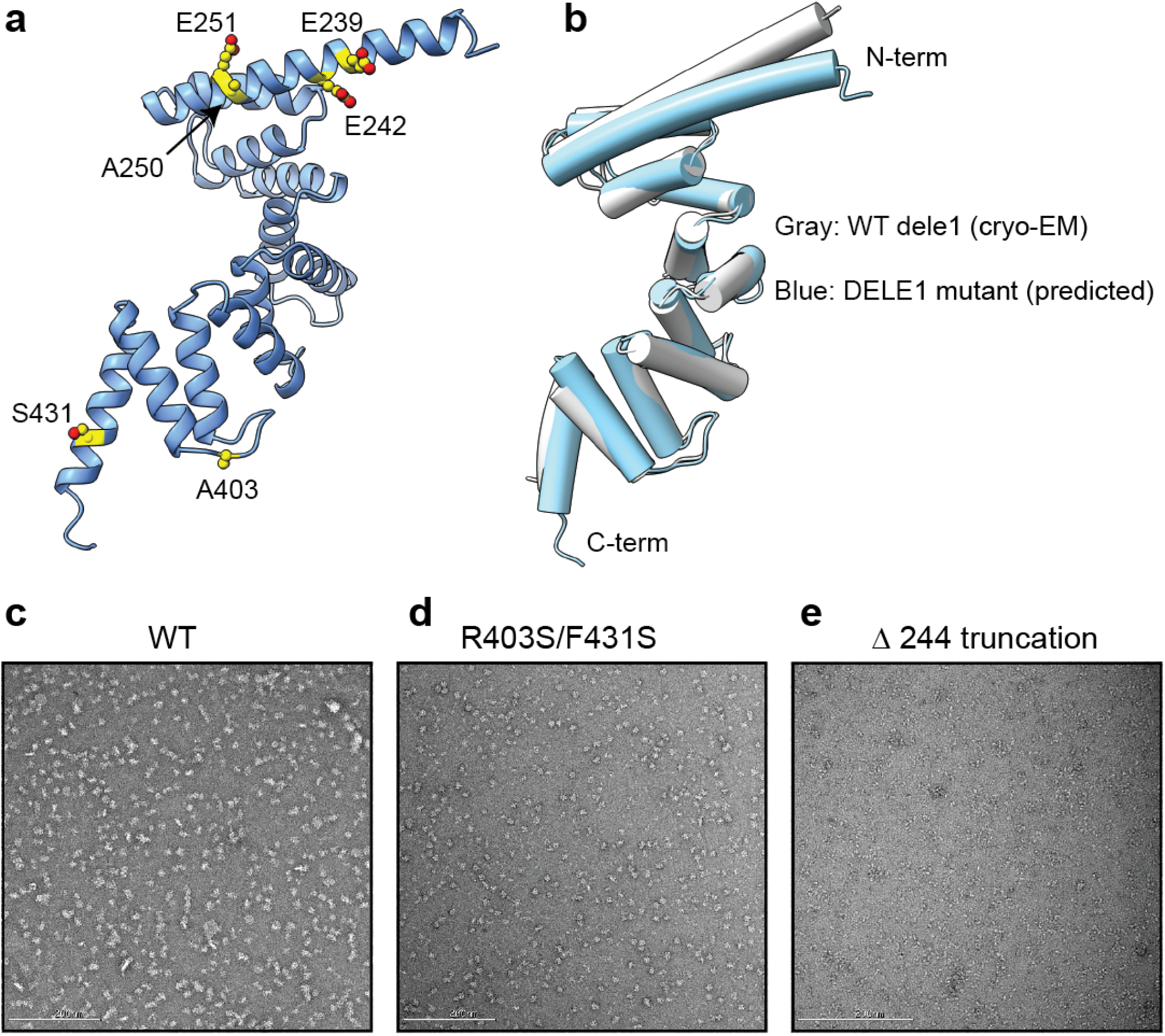
AlphaFold2-predicted models of DELE1 and negative stain EM. The AlphaFold2 predicted structure with the DELE1 CTD sequence harboring L239E, L242E, F250A, L251E, R403A, and F431S mutations. The mutated residues are denoted as yellow spheres. **(B)**Structural alignment of the DELE1 cryo-EM structure and AlphaFold2-predicted monomeric structure in (A), showing that the overall predicted structure of the mutant DELE1 is highly similar to the WT DELE1. **(C-E)** Representative negative stain EM images of the WT DELE1, the Interface II mutant (F431S/R403A), and the Interface I truncation (244).

**Supplementary Figure 4.**
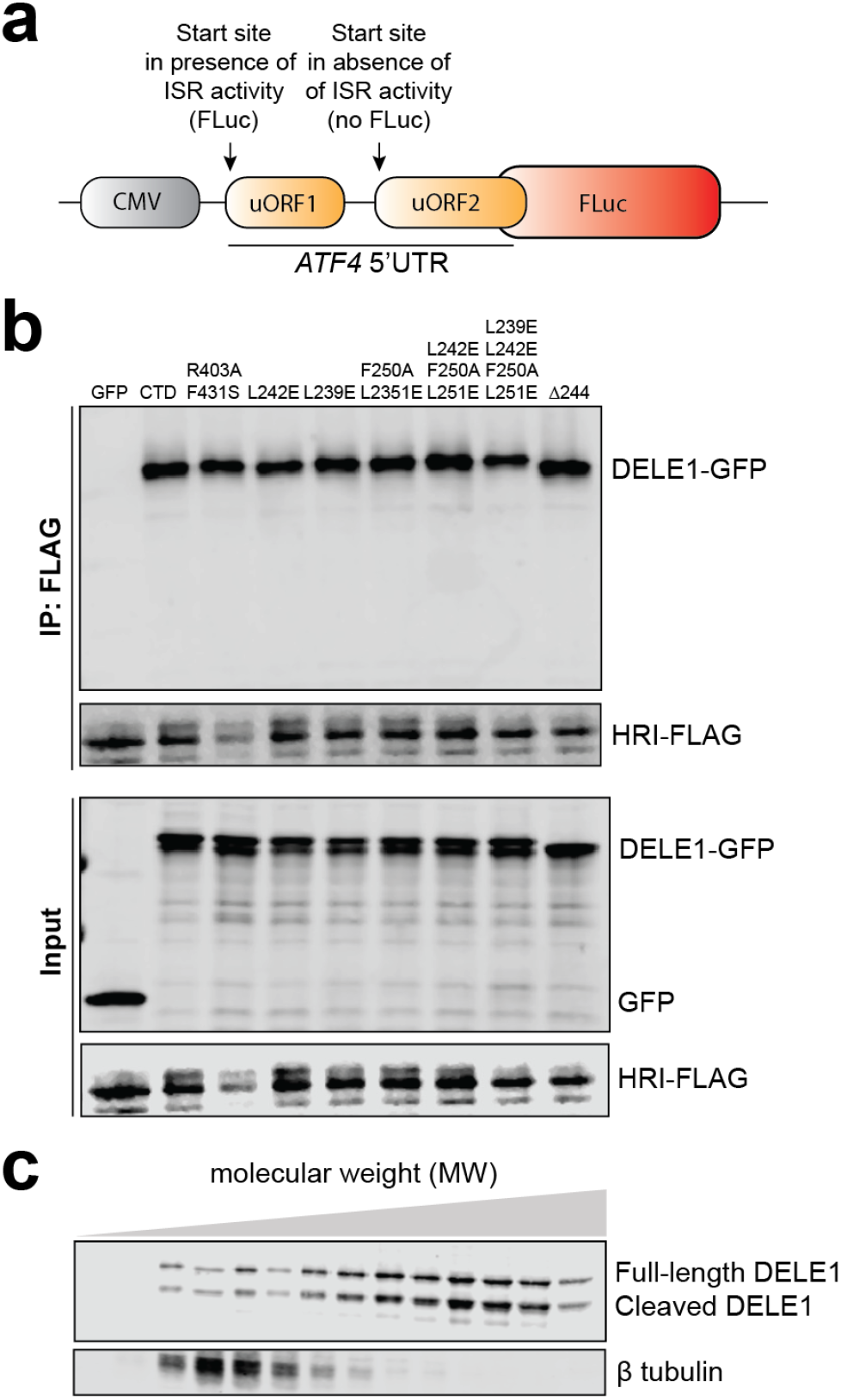
Interaction between HRI kinase and DELE1 mutants. **(A)** Description of the ATF4 reporter stable cell line, which contains the upstream open reading frames (uORF1 and uORF2) in the ATF4 5’ untranslated region (*ATF4 5’ UTR*). The ATF4 coding sequence was replaced by a luciferase gene (Fluc). **(B)** Co-immunoprecipitation of FLAG-tagged HRI and GFP tagged WT DELE1^CTD^ (CTD), and different DELE1 mutants. Anti-GFP antibody and Anti-FLAG antibody were used to detect DELE1 and HRI, respectively. **(C)** Immunoblot of the full-length DELE1-mClover protein from sucrose gradient fractions. The 14 sucrose gradient fractions from light to heavy molecular weights (left to right) were blotted with an anti-GFP antibody to detect the DELE1-mClover fusion protein. The upper and lower bands represent the full-length DELE1 and the cleaved DELE1, respectively. β tubulin was used as a control.

**Table S1.**
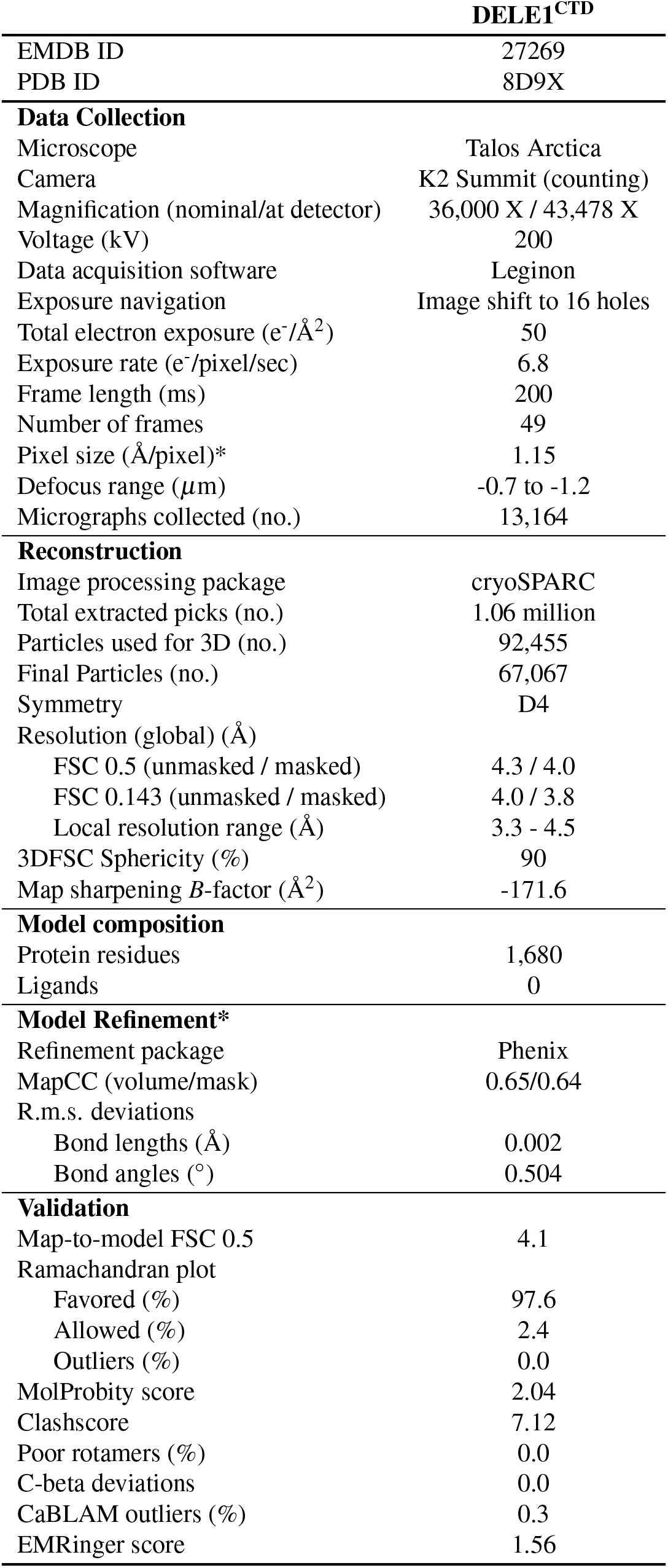
Cryo-EM data collection, refinement, and validation statistics.

